# Effect of salinity and water dilution on environmental DNA degradation in freshwater environments

**DOI:** 10.1101/2021.05.24.445344

**Authors:** Tatsuya Saito, Hideyuki Doi

## Abstract

Environmental DNA (eDNA) analysis methods have been developed to detect the distribution and abundance/biomass of organisms in various environments. eDNA generally degrades quickly, thus the study of eDNA degradation is critical for eDNA evaluation. However, there have only been a few studies of eDNA degradation experiments in which the salt concentration and water dilution were controlled. In this study, the effects of degradation were experimentally evaluated by controlling the salinity and water dilution of pond water. An experiment was conducted to evaluate the effects of salinity and dilution on eDNA detection with fragmental eDNA and free cell-derived eDNA using pond water, diluted pond water, and saline pond water. We quantified the eDNA copies of free cells, fragmental DNA, and the eDNA from *Cyprinus carpio*. In both the diluted and saline pond water, we found that the degradation rate of eDNA was much slower than that in pond water. Furthermore, the DNA concentration did not exponentially decrease in both the saline purified water and purified water samples. For the lower degradation rate in salt water, we interpreted that salts may affect DNA degradation factors such as microbe compositions and activities. The effect of salinity and dilution on eDNA detection provides fundamental information about the degradation process of eDNA, which is essential to understand the behavior of eDNA in natural environments.

## INTRODUCTION

Environmental DNA (eDNA) evaluation methods have been developed to monitor macroorganism communities and manage aquatic ecosystems [1–4]. eDNA is the DNA released by organisms into an environment, such as water or soil, and derives from the feces [5], skin cells [1], mucus [6], and secretions [7] of the organisms. In addition, this eDNA can be collected in aquatic systems [1,8]. The DNA sources are mainly fractions of cells or organelles but can also be free DNA fragments suspended in the water [9,10].

An understanding of eDNA degradation, which is a critical eDNA characteristic, is important for eDNA evaluation for both species distribution and abundance/biomass [11–13]. To reveal the states of eDNA, especially its degradation rate, many experiments have been conducted under various conditions [14,15], such as varied temperature [16,17], pH [15,18], and salinity [11].

eDNA is measured over an experimental period to evaluate eDNA release and degradation [14,15,19]. The degradation curves of the eDNA in most experiments have been observed to have exponentially declined [12] and eDNA concentrations can decay below the limit of detection in less than a week [13–15].

In previous meta-analyses of eDNA [12,20](, it was similarly shown that no significant difference could be observed in the eDNA degradation rates between freshwater and seawater. However, the results of degradation experiments using sea and pond water have shown that the eDNA degradation rate would be slower in the sea [13]. Collins et al. [11] found that there is a slower eDNA degradation rate in marine areas with higher salt concentrations. There have been only a few studies of eDNA degradation in which the salt concentration was controlled, despite its importance for eDNA evaluation in the field. Therefore, we evaluated the effect of salinity on eDNA degradation by conducting degradation experiments. Furthermore, water dilution can potentially reduce the factors of eDNA degradation, such as the enzymes and microbes that degrade DNA. Thus, we also tested the effect of water dilution on eDNA degradation in the same manner.

The aim of this study was to observe and compare the effects of salinity and water dilution on the eDNA degradation rate in freshwater environments. To understand the degradation in each DNA source, such as individual-derived, cell-derived, and fragmental DNA [13], we evaluated the effects of salinity and dilution on eDNA detection while considering the fragmental eDNA, free cell-derived eDNA, and eDNA derived from the resident species of the pond.

## MATERIALS AND METHODS

### Experimental design

We collected pond water, diluted pond water, and salined pond water and divided each water sample into bottles (Wide-Mouth Bottle, 500mL; AS ONE, Osaka, Japan) (Figure 1). We collected the pond water from an artificial pond in Kobe (the same pond used in Saito & Doi [13]). A solution of isolated cells (from *Oncorhynchus kisutch*) and fragmental DNA (from an internal positive control [IPC, 207-bp, 1.5 × 10^5^ copies; Nippon Gene, Tokyo, Japan]) was added to each bottle (Figure 1). The pond water contained the eDNA of the resident common carp (*Cyprinus carpio*). We used *O. kisutch* tissue for the isolation of cells because this species is not distributed in the pond. We conducted the experiment for seven days. Water samples (500 mL) from each bottle were filtered and collected using a Sterivex filter (0.45 µm pore size; Merck Millipore, Burlington, MA, USA; Figure 1). After extracting eDNA from the Sterivex filter, the copy number of each type of DNA contained in the Sterivex samples and filtrate was estimated by quantitative real-time PCR (qPCR, Figure 1).

**Fig 1.**
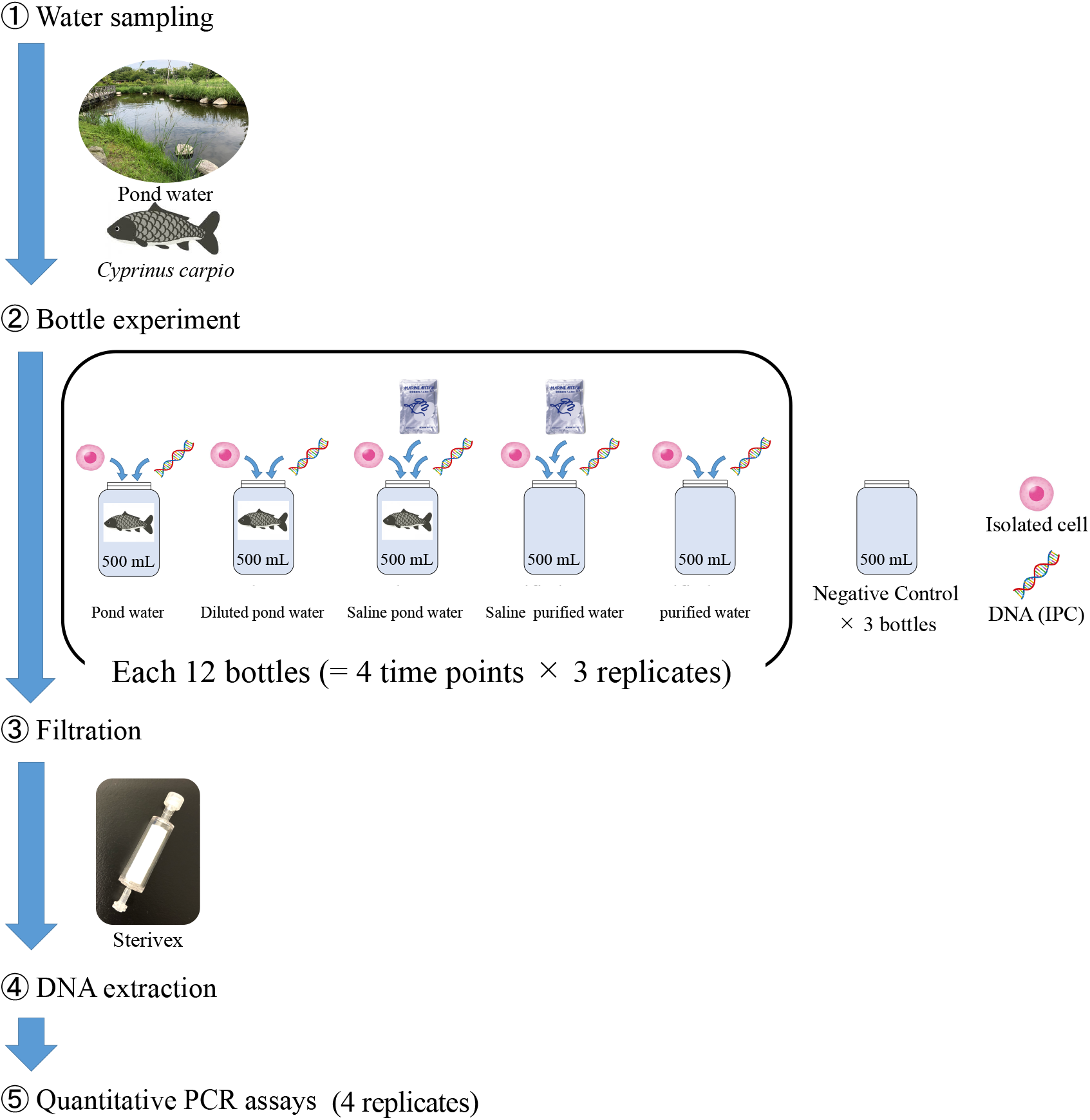
Experimental overview of the bottle experiments. We collected pond water, purified water, diluted pond water, saline pond water, and saline purified water and divided each type of water into 12 bottles. A solution of isolated cells (from *Oncorhynchus kisutch*) and fragmental DNA (IPC) was added to each bottle. The pond water was expected to contain the environmental DNA (eDNA) of *Cyprinus carpio*. We used *O. kisutch* tissue for the isolation of cells. We conducted the experiment for seven days. A Sterivex filter was used to filter 500-mL samples of water from each bottle. After extracting eDNA, the copy number of each type of DNA was estimated by quantitative real-time PCR.

### Bottle experiment

We collected the pond water from an artificial pond in Kobe, Japan (34º39’ 40” N, 135º 13’ 02” E) on July 16, 2020, using bleached tanks. We measured the salt concentration (salinity) and temperature of the collected water using a salinity meter (CD-4307SD; Mother Tool, Nagano, Japan) and a thermometer (ProODO; YSI, Tokyo, Japan), respectively. The salt concentration (salinity) and water temperature at the time of the water collection were 0.04 and 26.5 ºC, respectively.

For the saline water, artificial seawater powder (Marine Art BR; Osaka Yakken, Osaka, Japan) was added to the pond water and purified water to increase the salinity to 3.3, the mean seawater salinity around Japan. For the diluted pond water, the pond water and purified water (A300; AS ONE) were mixed at a ratio of 1:9. The pond water, purified water, diluted pond water, saline pond water, and saline purified water were each divided into 12 bottles (500 mL each). The bottles and equipment were sterilized with 10% commercial bleach (ca. 0.6% hypochlorous acid), (KAO, Tokyo, Japan) and washed with DNA-free distilled water to avoid DNA contamination.

Each bottle received 100 µL of a solution of isolated cells [1.0 × 10^5^ copies] and DNA (IPC). The bottles were incubated in the laboratory at about 25 ºC for a week. We collected and filtered 500 mL of the water from each bottle using 0.45-µm Sterivex filters (Merck Millipore) at 0, 3, 12, and 168 (day 7) h after the introduction of the cells and DNA. After filtration, approximately 2 mL of RNAlater (Thermo Fisher Scientific, Waltham, MA, USA) was injected into the Sterivex. As a filtration blank, the 500 mL of DNA-free water was filtered in the same manner after filtration of the samples to monitor cross-contamination. The Sterivex filters were immediately stored at –20 ºC until further analysis.

### DNA extraction

DNA was extracted from the Sterivex filter using a DNeasy Blood and Tissue Kit (Qiagen, Hilden, Germany) following Miya et al. [21] and Minamoto et al. [22]. The RNAlater was removed using a 50-mL syringe, and 440 µL of the mixture (220 µL of phosphate-buffered saline, 200 µL of Buffer AL, and 20 µL of proteinase K [Qiagen]) was added to the Sterivex filter. We incubated the filters on a rotary shaker (AS ONE) at 20 rpm for 20 min in a 56 ºC dry oven. We transferred the incubated mixture into a new 1.5 mL tube by centrifugation at 5000 g for 5 min. We then purified the mixture using a DNeasy Blood and Tissue Kit, and finally eluted the DNA in 100 µL of buffer AE from the kit. The extracted DNA from both methods was stored at –20 ºC until qPCR analysis.

### Quantitative PCR assays

We performed the qPCR analysis for *C. carpio* [2], *O. kisutch*, and the IPC [13]. We quantified the DNA concentrations by qPCR using a PikoReal™ qPCR system (Thermo Fisher Scientific). Each TaqMan reaction contained 900 nM of forward and reverse primers and 125 nM of a TaqMan probe in the 1× TaqPath™ qPCR master mix (Thermo Fisher Scientific). To this, 2 µL of the sample template was added to reach a final volume of 10 µL. A four step dilution series containing 1.5 × 10^1^ to 1.5 × 10^4^ copies was prepared and used as quantification standards. For the standard curves, we used target DNA cloned into a plasmid.

A quantitative PCR was performed with the following conditions: 2 min at 50 ºC, 10 min at 95 ºC, and 55 cycles of 15 s at 95 ºC and 1 min at 60 ºC. Four replicates were performed for each sample, and four replicate negative non-template controls (NTC) containing DNA-free water instead of template DNA were included in all PCR plates. We performed the qPCR procedures according to the MIQE checklist [23]. The PCR and qPCR were set up in two separate rooms to avoid DNA contamination.

The qPCR results were analyzed using PikoReal software ver. 2.2.248.601 (Thermo Fisher Scientific). The R^2^ values of the standard curves ranged from 0.985– 0.998 (Supplementary Information) and the PCR varied from 91.07–101.68%. The concentration of DNA in the water collected (DNA copies mL^−1^) was calculated based on the volume of filtered water. DNA copy numbers were evaluated including negative amplifications set as zero values. In our previous study [13], we have already performed a limit of detection (LOD) test for the PCR assay, which resulted in one copy for the LOD.

### Statistical analysis

Statistical analysis and data plotting were performed using R software version 3.6.0 [24]. We used the Single First-Order rate model (SFO) as the degradation model because the SFO was the most effective model of degradation in Saito and Doi [13]. The SFO establishes a simple procedure for determining a first-order rate constant from the degradation. The model equation is as follows:

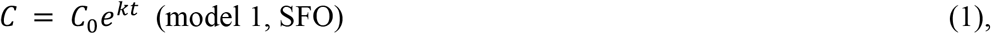

where *C* is the eDNA concentration at time t, *C*_0_ is the eDNA concentration at time 0 (i.e., the initial eDNA concentration), and k is the degradation rate constant per hour. We performed modeling using the “mkin” package version 0.9.49.8 in the R software. We evaluated the fit of the models using the chi-squared error level [25](Boesten et al.,2005). Significant differences in the model coefficients were evaluated by overlapping the 95% confidential intervals (CIs) of the coefficients (i.e., α = 0.05).

## RESULTS

### Degradation of eDNA in saline pond water

We detected all the targeted DNA of *C. carpio*, the *O. kisutch* cells, and the IPC using qPCR in saline pond water (Figure 2). The degradation rates in the saline pond water were significantly lower than those in the regular pond water for all three DNA sources (Table 1, Figure 3). We could not detect the eDNA of *C. carpio*, the *O. kisutch* cells, or the IPC on day seven in the pond water. However, we detected the eDNA of *C. carpio* and the IPC in the saline pond water up to day seven. There were no amplifications from the filter, extraction blanks, and NTCs in this experiment or in the following experiments.

**Table 1.** Degradation rate constant of Single-First Order models. The values are slope k and the values in parentheses are the lower and upper confidential intervals of slope k.

| Water | IPC | Okis | Cyca |
| --- | --- | --- | --- |
| Pond | 0.293 (0.2202, 0.3899) | 0.4510 (0.3612, 0.5631) | 0.1107 (0.0813, 0.1506) |
| Saline pond | 0.00757 (0.002361, 0.024270) | 0.01746 (0.00659, 0.04626) | 0.01465 (0.00344, 0.06244) |
| Diluted pond | 0.009872 (0.004441, 0.021950) | 0.03306 (0.02026, 0.05394) | 0.02519 (0.01592, 0.03986) |

Cyca, *Cyprinus carpio*; IPC, internal positive control; Okis, *Oncorhynchus kisutch* cells.

**Fig 2.**
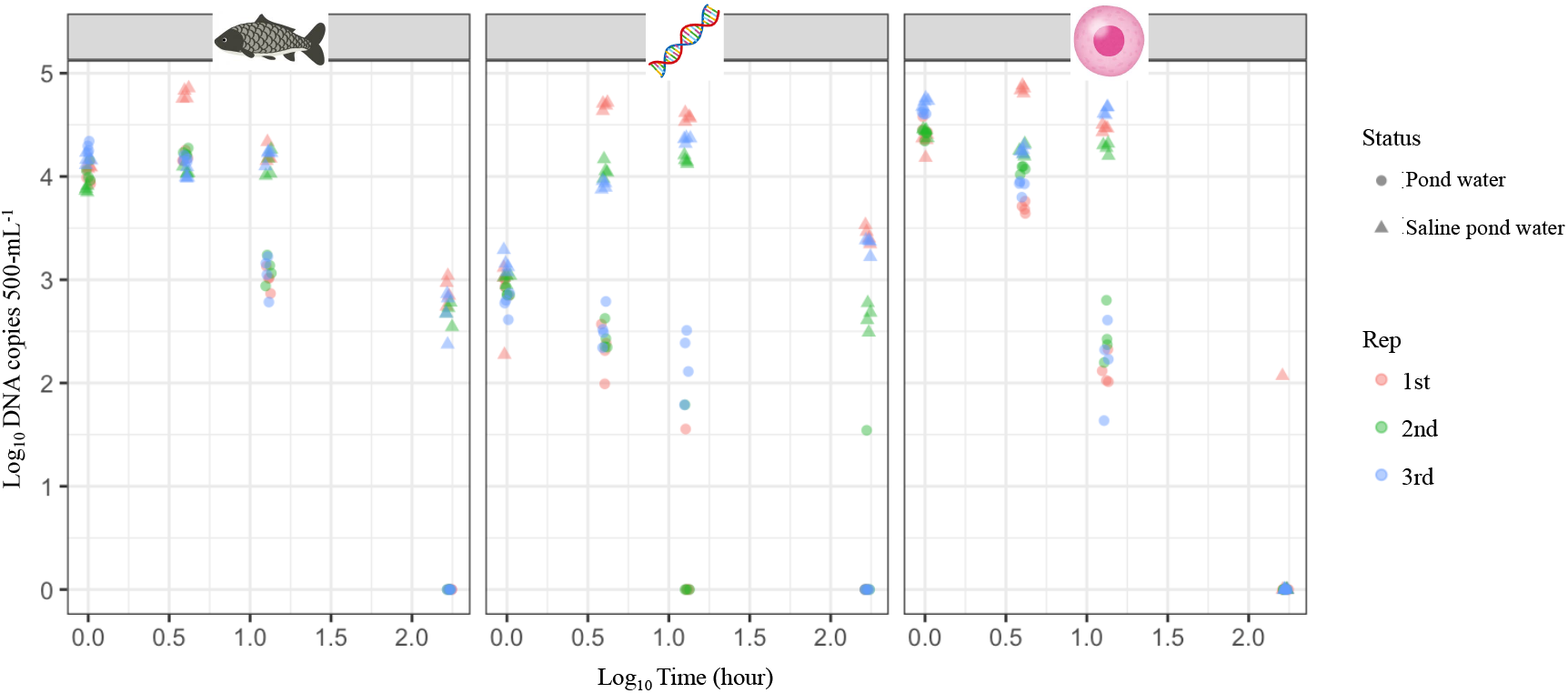
Relationship between the environmental DNA (eDNA) concentrations of the water source (pond water and saline pond water). The dots indicate the eDNA concentrations of the targets at each time point under two water conditions: pond water, circles; saline pond water, triangles (N = 12 for each time point: 1st replication, red; 2nd replication, green; 3rd replication, blue).

**Figure 3.**
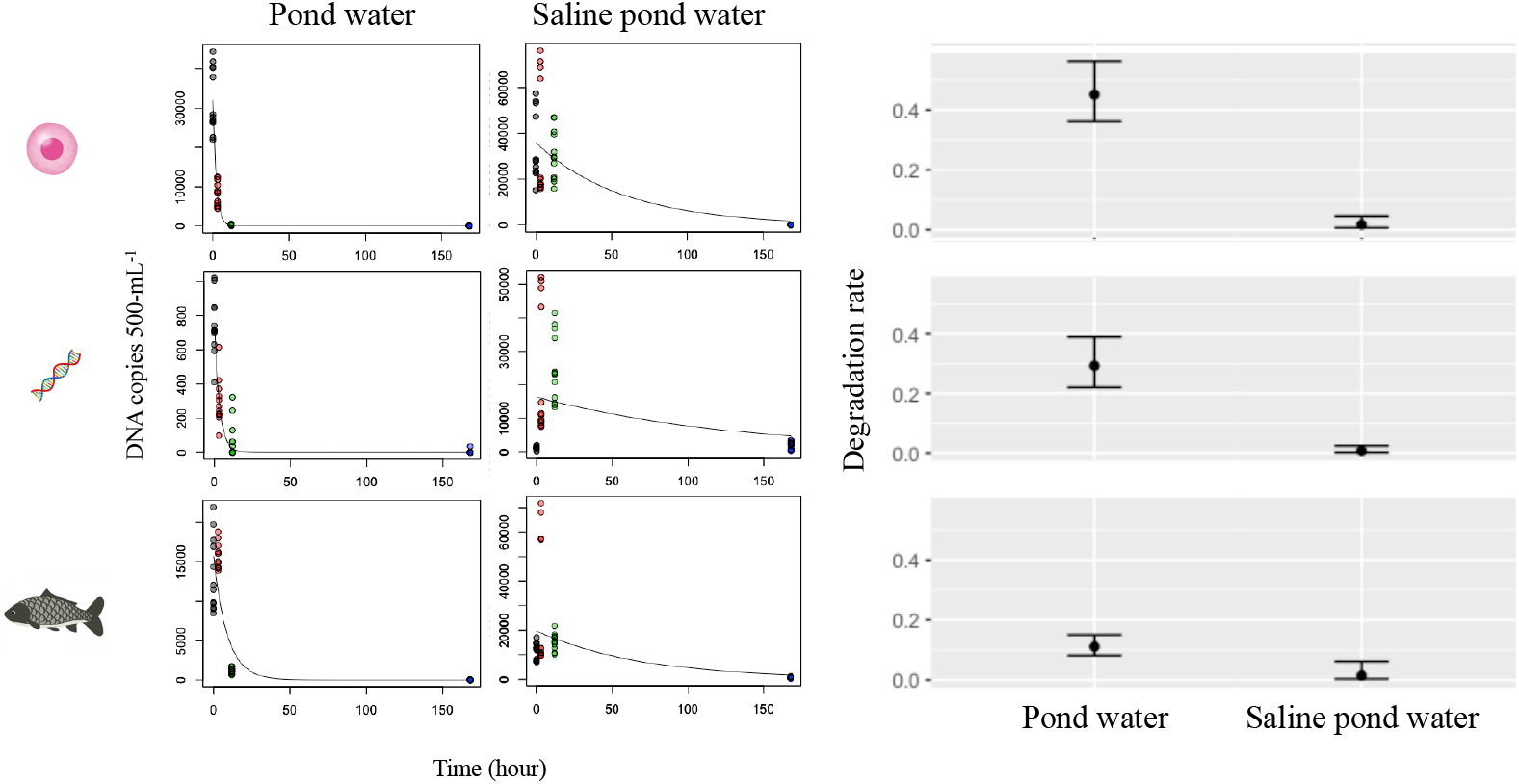
Degradation curves of the Single-First Order (SFO) model and the rate constant for the bottle experiments in the saline pond water and pond water. The dots indicate the environmental DNA concentrations of the targets (*Cyprinus carpio*, the internal positive control [IPC], and *Oncorhynchus kisutch* cells) at each time point with different colors (N = 12 for each time point). The left degradation curves show each target (*O. kisutch* cells, the IPC, and *C. carpio*) in the pond samples. The right decay curves show each target (*O. kisutch* cells, the IPC, and *C. carpio*) in the saline pond samples. In the right-hand plots, the slopes (k) of each target (*O. kisutch* cells, the IPC, and *C. carpio*) are shown with 95% confidential intervals.

The degradation rate constant (k) of the cells, IPC, and *C. carpio* were significantly different between the saline pond and pond samples when comparing the 95% CIs (Figures 2 and 3). The degradation rates of the saline pond were significantly lower than those of the pond water for all three DNA sources.

### Degradation of eDNA in saline purified water

We detected all the targeted DNA of the *O. kisutch* cells and the IPC using qPCR in the saline purified water (Figure 4). We detected the *O. kisutch* cells and the IPC DNA in the purified water and saline purified water, respectively, up to 168 h. The DNA concentrations of the cells and IPC DNA did not decrease exponentially after they were added (0 h).

**Fig 4.**
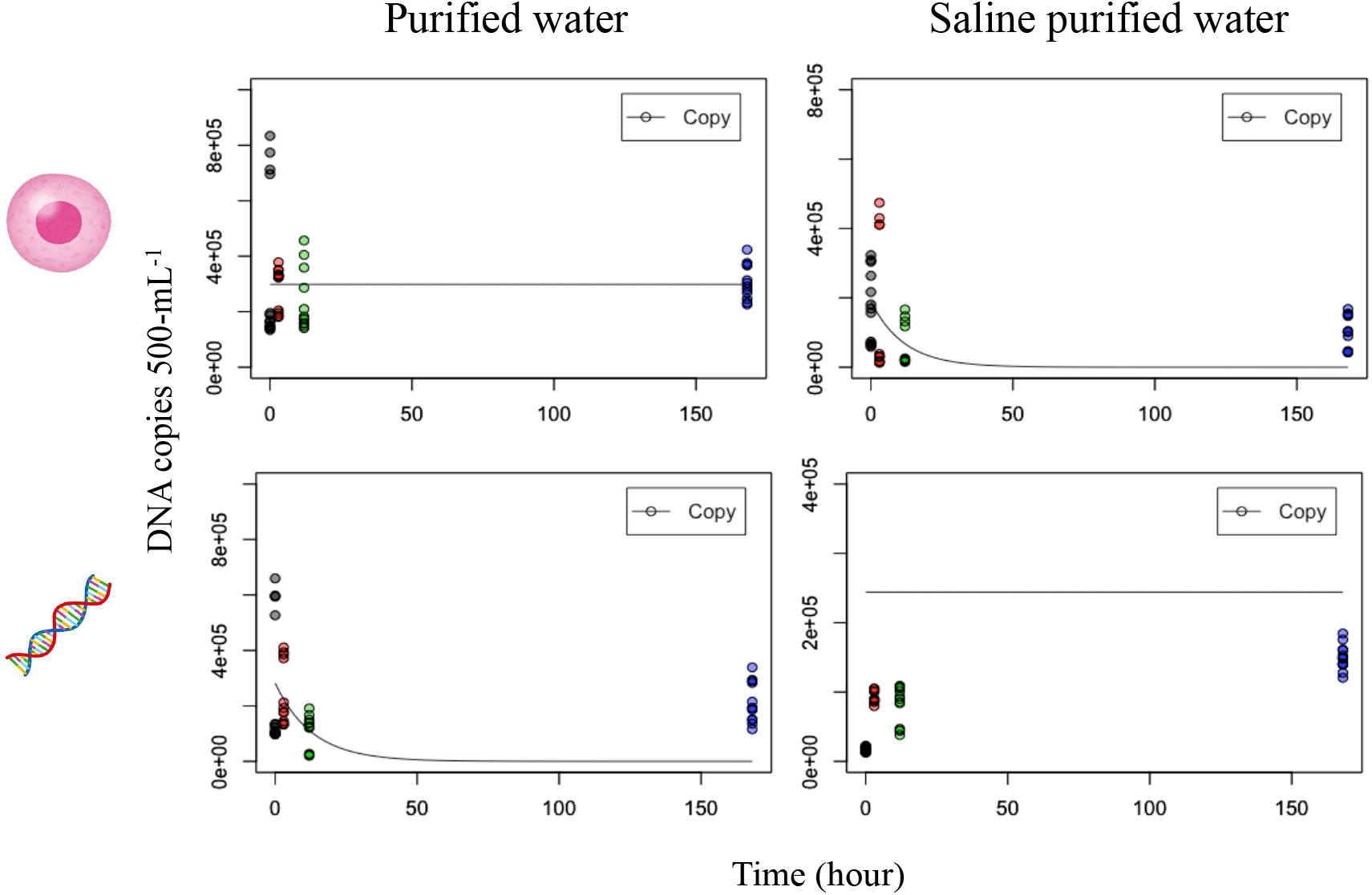
Degradation curves of the Single-First Order model (SFO) for the bottle experiments in the saline purified water and purified water. The dots indicate the environmental DNA concentrations of the targets (the internal positive control [IPC] and *Oncorhynchus kisutch* cells) at each time point with different colors (N = 12 for each time point). The left degradation curves show each target (*O. kisutch* cells and the IPC) in the purified water samples. The right decay curves show each target (*O. kisutch* cells and the IPC) in the saline purified water samples.

### Degradation of eDNA in diluted pond water

We detected three of the targeted DNA types, *C. carpio, O. kisutch* DNA from the cells, and the IPC, using qPCR in diluted pond water (Figure 5). The degradation rates of the diluted pond were significantly lower than those of the pond water for all three DNA sources (Table 1, Figure 6). We could not detect the eDNA of *C. carpio*, the *O. kisutch* cells, and the IPC at 168 h in the pond water. However, we detected the eDNA of *C. carpio* and the IPC in the diluted pond water up to 168 h.

**Fig 5.**
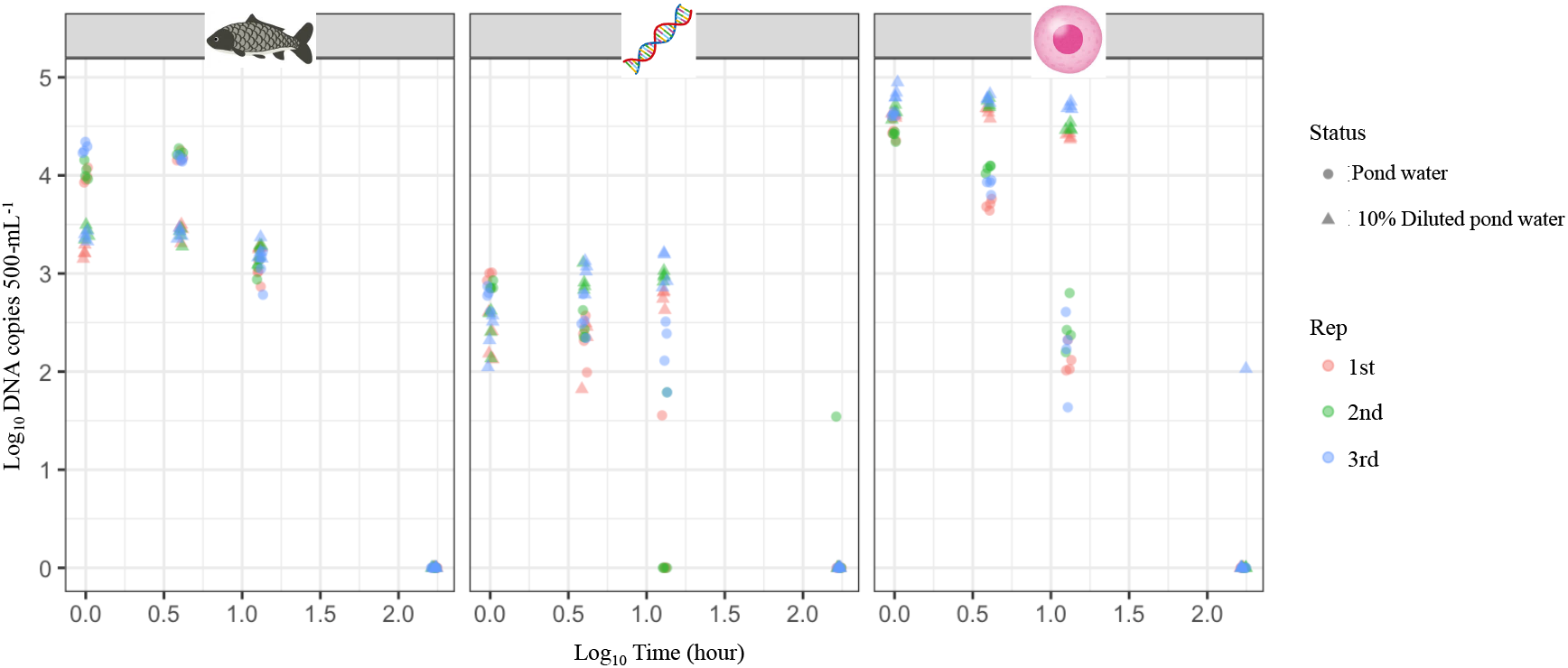
Relationship between the environmental DNA (eDNA) concentrations of the water source (pond water and 10% diluted pond water). The dots indicate the eDNA concentrations of the targets at each time point under two water conditions: pond water, circles; diluted pond water, triangles (N = 12 for each time point: 1st replication, red; 2nd replication, green; 3rd replication, blue).

**Fig 6.**
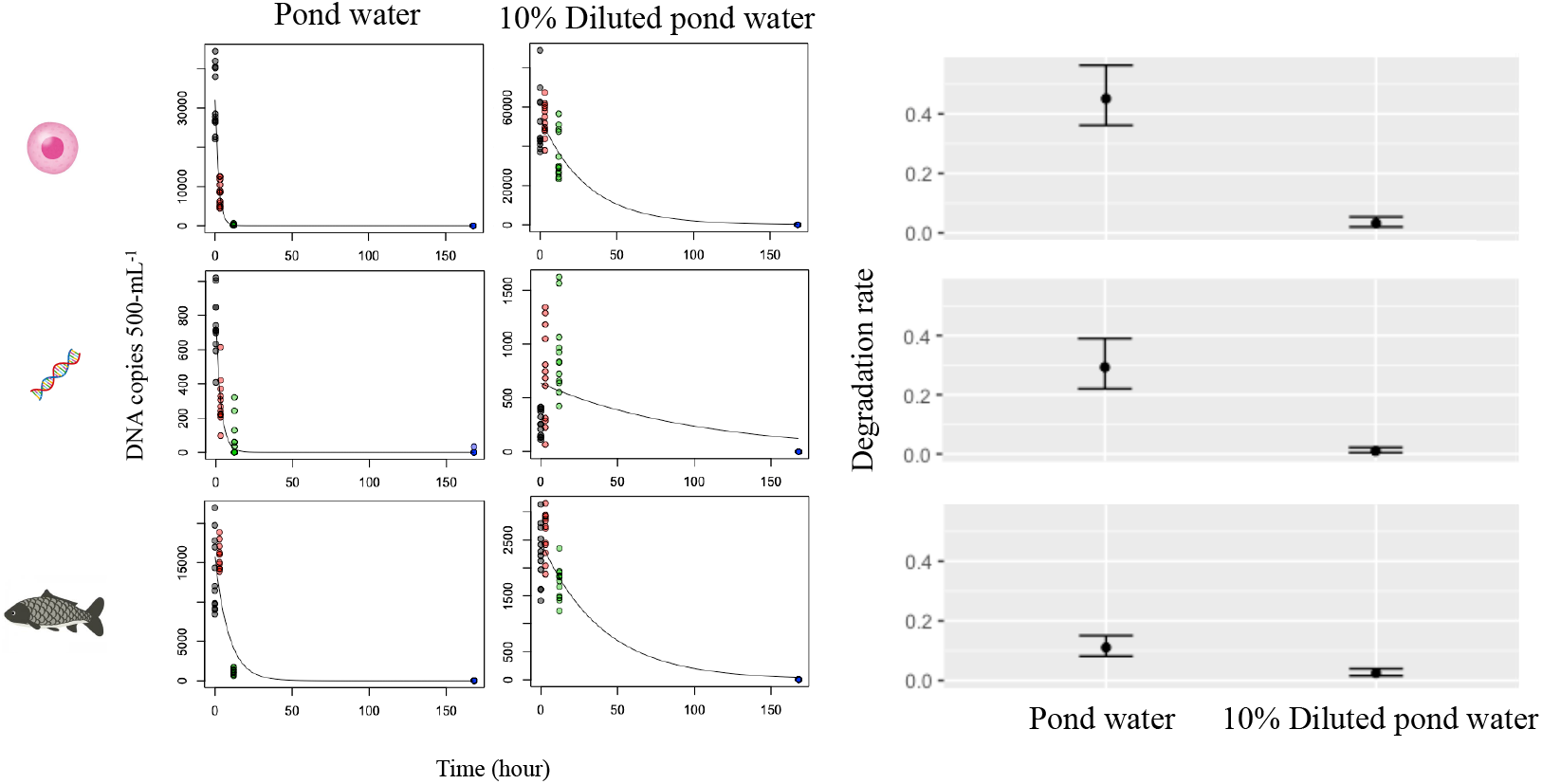
Degradation curves of the Single-First Order model (SFO) and the rate constant for the bottle experiments in diluted pond water and pond water. The dots indicate the environmental DNA concentrations of the targets (*Cyprinus carpio*, the internal positive control [IPC], and *Oncorhynchus kisutch* cells) at each time point with different colors (N = 12 for each time point). The left degradation curves show each target (*O. kisutch* cells, the IPC, and *C. carpio*) in the pond samples. The right decay curves show each target (*O. kisutch* cells, the IPC, and *C. carpio*) in the diluted pond samples. In the right-hand plots, the slopes (k) of each target (*O. kisutch* cells, the IPC, and *C. carpio*) are shown with 95% confidential intervals.

The degradation rate constant (k) of the cells, IPC, and *C. carpio* were significantly different between the diluted pond and pond samples when comparing the 95% CIs (Figure 6). The degradation rates in the diluted pond water were significantly lower than those in the pond water for all three DNA sources.

## DISCUSSION

We found that the DNA concentrations of the *C. carpio, O. kisutch* cells, and the IPC did not decline exponentially in both the saline purified water and purified water samples. Our present results supported our previous study [13]. Furthermore, we detected the DNA in the saline purified water. This result showed that the increased salinity in the saline sample did not have any effect on DNA detection.

The degradation rates in the saline pond samples were significantly lower than those in the pond water for all three DNA sources. This result might suggest that salinity suppresses the degradation of eDNA. However, the results of a meta-analysis of eDNA degradation showed that the eDNA degradation rates between freshwater and seawater are not significantly different [12,20]. A previous study [11] found salinity to be a better predictor of eDNA decay than pH, with salinity varying more between locations. They also showed that salinity itself may not be entirely responsible for the difference in degradation rate, but rather that it is associated with abundances or communities of microbes. In fact, the characteristics of microorganisms involved in DNA degradation may vary depending on the environment. For example, microorganisms living in freshwater are susceptible to salinity and cannot adapt to the rapid environmental change caused by salt addition, which is thought to reduce their activity and suppress DNA degradation. Therefore, the results of our study suggest that salts would be unlikely to protect DNA and may affect DNA degradation factors, including microbe composition changes and the activity of DNA enzymes in the water. Further evaluation of microbe compositions and DNA enzymes in salt water is needed to gain a deeper understanding of the degradation of eDNA.

The degradation rates in the diluted pond samples were significantly lower than those in the pond water for all three DNA sources. In the pond water diluted 10 times, the initial DNA concentration was reduced to one-tenth, but the degradation rate was slower than that in the pond water. This is thought to be due to the dilution of the degradation factors as well as the eDNA. Takasaki et al. [26] showed that pre-filters that remove humic substances such as humic acid and fulvic acid are effective in detecting eDNA. Our results showed that simply diluting eDNA without removing its degraders and inhibitors was effective. However, the DNA was not detected after seven days. Therefore, it was found that even if the concentration of the degradation factor is low, the degradation progresses over time.

Our experiments provide new findings on eDNA degradation; however, there were some limitations owing to the experimental design. First, we performed the experiment using only one site for collecting the pond samples. Therefore, it is unclear whether similar DNA degradation rates exist seasonally and in the other aquatic habitats such as river and wetlands. Experiments using a selection of site replicates from various habitats need to be performed to achieve a more generalized understanding of eDNA degradation. The evaluation of eDNA degradation while comparing different environmental conditions (e.g., salinity, water temperature, pH, chlorophyll, and microorganism population) may reveal what is affecting eDNA degradation in general.

In conclusion, we found that the DNA concentrations of the *C. carpio, O. kisutch* cells, and the IPC did not decline exponentially in both the saline purified water and purified water samples. The degradation rates of the saline pond and pond samples were significantly different. The degradation rates of the diluted pond and pond samples were significantly different. A greater understanding of and the accumulation of basic information about eDNA would improve eDNA analysis methods and enable researchers to maximize the potential of future eDNA methods.

## ACKNOWLEDGMENTS

This study was supported by the Environment Research and Technology Development Fund (JPMEERF20164002 and JPMEERF20204004) of the Environmental Restoration and Conservation Agency, Japan.

## DATA ACCESSIBILITY

All data, including the raw values for the qPCR experiments, are included in the supporting information.

## Notes

### Competing Interest Statement

The authors have declared no competing interest.

